# Evolution of MEG: a first MEG-feasible fluxgate magnetometer

**DOI:** 10.1101/2021.03.22.435762

**Authors:** N. Koshev, A. Butorina, E. Skidchenko, A. Kuzmichev, A. Ossadtchi, M. Ostras, M. Fedorov, P. Vetoshko

## Abstract

In the current article we present a first solid-state sensor feasible for magnetoencephalography (MEG), and working at room temperature. The sensor is a fluxgate magnetometer based on yttrium-iron garnet films (YIGM). In this feasibility study we prove the concept of usage the YIGM in terms of MEG by registering a simple brain induced field: the human alpha rhythm ^1^. All the experiments and results are validated with usage of another kind of high-sensitive magnetometers - optically pumped magnetometer (OPM), which currently appears to be well-established in terms of MEG.

## 1 Introduction

Magnetoencephalography (MEG) is an attractive neuroimaging modality that combines non-invasiveness with high spatial and temporal resolution. These properties place MEG among the most informative neuroimaging tools capable of localizing neuronal activity with distinct temporal structure and suitable for studying complex functional integration processes. The first demonstration of MEG in humans dates back to 1972 when D.Cohen [1] used Superconducting Quantum Interference Devices (SQUIDs) to register human alpha activity. Since then MEG found numerous applications in both medicine [2–4] and neuroscience [5–8].

Currently, SQUID based MEG device (SQUID-MEG) implemented in the form of a fixed-size helmet-dewar remain the most widely used MEG instrument. While SQUID-MEG has been successfully utilized in experimental and clinical neurology the technology behind SQUID-based MEG systems limits the range of exciting applications. The system is costly to purchase and maintain due to the constant need for cooling the sensors in the liquid helium. In addition, the application of such a system is difficult for subjects with small heads (babies, children). For adult subjects, the distance between the sensors and the human head is about 2-3 cm (the thickness of the dewar walls) and noticeably increases for children, severely limiting the sensitivity of the device due to the attenuation of the magnetic field in inverse proportion to the square of the distance between the current source and the sensor [9]. Despite all the advantages and uniqueness of the spatial and temporal resolving properties the listed technological shortcomings precluded a wide spread of MEG neuroimaging modality currently available only at several hundred locations worldwide. Recent technological developments in the field of atomic magnetometry hold promise of lifting several limitations inherent to the current MEG systems.

The first demonstration of a non-cryogenic sensor with a SQUID-system level sensitivity 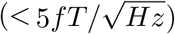 was described by Schwindt et al. at 2010 [9] who used optically pumped magnetometers (OPM) to register brain activity. Optically pumped magnetometers (OPM) operate in the so-called spin-exchange relaxation free (SERF) mode [10]. The OPM is a compact device consisting mainly of a laser, a photodiode, and a gas cell. The gas cell is a sensitive element of the OPM and may contain atoms of one of the alkali metals: Cs, Rb, and others. In SERF mode, magnetometers operate at a high alkali metal vapor density and near-zero background magnetic field, which allows OPM to achieve a sensitivity level comparable to SQUID systems with theoretical limit of 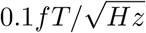 for compact implementations.

Unlike SQUID, OPMs do not require cooling to cryogenic temperatures for operation. They can be assembled into flexible arrays around the head, adapting to any head size and shape. This way the sensitive element of the atomic magnetometers appear located closer to brain sources because of only a 6 *−* 6.5 mm distance between the vapor cell and the outer housing of the sensor. In addition, OPM showed greater robustness in experiments compared to SQUID ([9, 11–13]). The 20-channel system was able to achieve high quality response recording with standard peak recognition while showing semi-quantitative similarity with SQUID recordings [14]. Nowadays, the OP-MEG studies are being continued and there are several OP-MEG systems in the world containing up to 50 channels [15].

While OPM MEG systems represent a very significant step forward and hold promise to revolutionize the entire neuroimaging field the state-of-the-art, OPM sensors have significant disadvantages such as low dynamic range and bandwidth, high temperature of operation and non-solid state nature of the device. Here we for the first time propose a solid state sensor working at room temperature and usable for MEG: a high-sensitivity flux-gate magnetometer based on Yttrium-Iron Garnet Films (further - YIGM). The proposed sensor has a range of serious advantages over both SQUID and OPM based solutions.

Indeed, SQUID sensors require to be placed into a dewar with 2.5-3 cm thick walls while OPMs sensitive element is a vapor cell heated to around 100 C which requires insulation and results in 0.6 cm separation between the proximal wall of the vapor cell and the scalp. Unlike these two sensors, the proposed solid state YIGM sensor is flat by design (see below in Section 2) and operates at the room temperature. This makes it potentially possible to place the YIGM sensitive element at less that 2 mm away from the scalp surface when registering tangential components of the magnetic field.

Thus, since the magnetic field from a dipolar source decreases proportionally to the square of the distance, YIGM located closer to the sources can register deeper and/or weaker brain electrical activity as compared to the SQUID or OPM sensors. Moreover, the theoretical limit of YIGM sensitivity is about 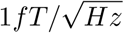 [16], which is comparable or higher to that of the currently implemented OPM devices.

Another set of serious advantages as compared to OPM is related to SERF-free nature of the YIGM sensor. Usage of zero-field resonance imposes serious limitations on ambient magnetic fields and its dynamics. For example, the dynamic range of the Quspin QZFM Gen. 2.0 is *±*5*nT* (see, e.g., site Quspin.com). As opposed to that, the dynamic range of currently used YIGM is 10*µT*, and further can be improved up to 100*µT* [17]. Being a SERF-free device, the YIGM required to be calibrated only once. There is no need for recalibration due to strong magnetic contaminations like opening the door of magnetically shielded room, etc. Moreover, theoretically, unlike current OPM devices operating in the SERF mode and requiring low (*<* 50*nT*, [18]) ambient fields achieved with costly magnetic shielding the YIGM sensor with its large dynamic range can potentially be used with less magnetic shielding.

The power consumption of OPM is 4.5W, about 700mW of which are irradiated by a sensor head itself [18]. Thus, the helmet containing, e.g., 50 OPMs, will have power consumption of about 35-40 W. On the other hand, the power consumption of the whole human body is about 100 W [19]. Therefore we can expect some discomfort and need for cooling in the OPM-based MEG-systems. In contrast the YIGMs operate at room temperature and have power consumption is about 100mW [17] which avoids any problems with overheating and the associated discomfort of the subject under the measurement. Lastly, due to simplicity of construction, YIGMs are potentially cheaper as compared to both OPMs and SQUIDs. YIGM does not contain parts with limited lifetime, which makes it durable and guarantees no performance deterioration.

Our aim here is to present the initial feasibility study of the YIGM sensors in MEG applications by registration of human alpha rhythm. In order to validate our measurements, we use the OP-MEG system, which has been chosen due to comparable to YIGM scalp-sensor distances.

## 2 Description of the sensor

The core of the YIGM sensitive element is a single-crystal ferrite-garnet epitaxial film shaped in a special way. Operation principle of this device related to the fluxgate type of the sensors with saturated core. Saturation in our sensor provided by rotation of magnetization in a circular rotating magnetic field. Theoretically it was predicted that this technology has a potential to increase its performance to 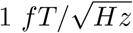 [16].

In our experiments we managed to obtain an unprecedented level of the noise power spectral density for the fluxgate type solid-state sensors due to the unique properties of ferrite-garnet core and a thorough preparation of our measurement setup. In order to conduct MEG experiments the sensor was fixed on a massive wooden stand. The stand is placed on a damping material made of soft polymer to prevent mechanical vibrations (see Figs. 1, a) and 3).

**Figure 1.**
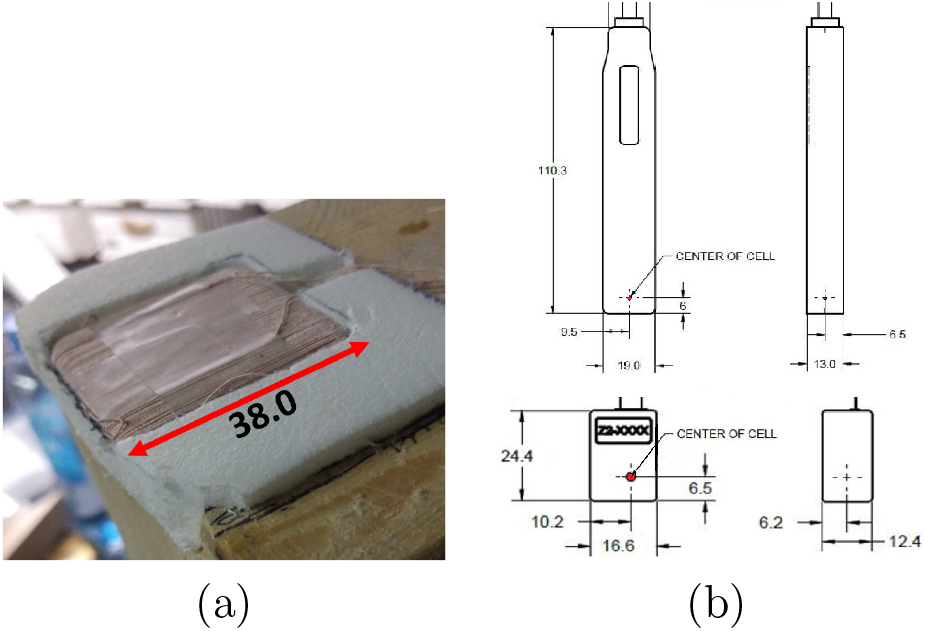
Sensors’ linear sizes and sensitive axes, where “T” stands for tangential, while “N” - for normal field component: a) YIGM sensitive element in winding; b) Sensor head of OPM QZFM Gen 1.0 (top) and 2.0 (bottom) magnetometers. Source: Quspin.com.

The linear dimensions and the arrangement of the sensitive axes of the YIGM sensor and the QuSpin OPM QZFM Gen-2 device can be compared in Fig.1, a) and b) correspondingly. The film represents a 40 *×* 40 mm patch with thickness of 1 mm (Fig. 1a). Taking into account thickness of the winding (1 mm per side), the overall size of the sensitive sensor head is about 3 mm. Depending on the relative position with respect to the subject’s head the sensor can detect either two tangential or one normal and one tangential components of the magnetic field as explained later in Section 3.6.

### 3 Methodology

In our feasibility study presented here we have chosen to perform comparative analysis of human alpha rhythm registered with OPM and YIGM sensors. The existence of a clear behavioral correlate, high signal amplitude and comparative simplicity of the alpha-wave registration experiments make this rhythm a good candidate for the proof-of-concept study illustrating the feasibility of YIGM sensors for registration of brain activity. One of the most attractive features of the occipital alpha rhythm is its connection to the eye-closed and eyes-open state of a subject which can be used to establish the physiological relevance of the obtained measurements.

#### 3.1 Alpha wave

Cortical encephalographic alpha rhythm is generally regarded as the electrophysiological correlate of the awake but resting conscious state. It is the most prominent signal of the ongoing EEG/MEG from wake human subjects at rest. In the last years, the reciprocal interaction between the lateral thalamic nuclei (specific relay nuclei) and the nucleus reticularis of the thalamus has been proposed to represent the central process in modulating cortical alpha activity [20], [21]. P.Roland [22] estimated the size of activated cortical areas covering comparatively big area (more than 6 cm^2^). The traditionally defined frequency range of the alpha band is (8, 12) Hz. However, individual variations are quite large, and the mean frequency of alpha varies as a function of age, gender, and even intelligence. It is quite low in frequency in the infant human (*<* 7 Hz), reaches its maximum in young adulthood, and declines with age [23]. Relative power of the alpha band significantly lower before detected stimuli in line with significantly higher amplitudes of the ERPs in a task of stimulus detection where 50% of the stimuli undetectable [24].

Sources of alpha rhythms may be found concentrated mainly in the region around the Calcarine sulcus, with most sources located within 2 cm of the midline [25]. The sources produce magnetic fields with magnitude of hundreds fT while registering with conventional SQUID-MEG. The magnitude of the alpha-induced magnetic field may reach 0.1-1 pT in dependence on experiment conditions and source-sensor distance. The first registration of magnetic field related to alpha-rhythm has been provided by David Cohen [1].

#### 3.2 General description of the experiments

The first-time application of a new kind of sensor requires a sophisticated validation of the obtained results. The laboratory use of the OPM-based MEG systems is well established by now. Given that the OPM sensors can be positioned with respect to the head similarly to the proposed YIGM sensor, we used OPM based alpha-band measurement for validation of the results obtained with YIGM. We performed our experiment on the group of three healthy adults^2^ (below - subjects) which further contributes to the reliability of our conclusions.

The conducted experiments can be divided into several parts described in details below. We start with the calibration of the YIGM sensor by registration of magnetic induction of known magnitude followed by the empty room noise spectrum recordings performed one after another with YIGM and the OPM sensors placed in the exact same place in the MSR. This allows us to compare the spectral noise profiles registered at the same location with appropriately oriented OPM and YIGM, to explore the sensor noise and distinguish it from the ambient residual field present in the MSR.

After that we use an array of the OPMs and individually for each subject find the OPM position with the largest magnitude of the occipital alpha rhythm observed in the eye-closed condition. We then place our YIGM at these locations and conduct the main experiment of the current article demonstrating the feasibility of registering human occipital alpha waves with solid-state YIGM sensors.

#### 3.3 The calibration of YIGM

We start with YIGM calibration process by registering a known magnetic field at the axis of the test coil with a weak current. In order to minimize the influence of the external fields on the calibration procedure the test coil and the YIGM sensor were both placed inside the demagnetised 3-layer permalloy schield. A 120 mm in diameter test coil comprising a single turn of the wire was connected through 240 kOmh of load resistance to the signal wave generator producing 10 mV RMS voltage at 10 Hz. Therefore, it can be concluded that the magnetic field in the center of the coil is approximately 600 fT.

#### 3.4 Empty room study

The experiment pursues two main goals. The first one is to determine the sweet-spot in the MSR with the minimum ambient noise and to ensure that the level of noise is suitable for our alpha rhythm registration experiments. The second goal is to study the intrinsic noise properties of our YIG sensor. Using the OPM sensors we sampled the ambient field inside the MSR and found a sweetspot, see Results section for the exact values of the observed fields.

The recordings of the background magnetic field inside MSR were performed one after another with YIGM and OPM sensors placed at the discovered sweet-spot location with minimal magnetic interference. The orientation of sensor sensitive axes in both cases was identical. All subsequent measurements (registration of alpha waves) were performed at the same location with minimal interference and with the same orientation of YIGM.

#### 3.5 The OP-MEG alpha activity registration experiment

The location and most notably orientation of an equivalent current dipole approximating the generator of the occipital alpha activity vary significantly across individuals. Therefore, we may expect that the maximum magnitude of the normal and tangential components of the magnetic field will be achieved at different locations in different subjects. Therefore, we start with registration of alpha waves using the OPMs QuSpin Zero Field Magnetometer (QZFM, QuSpin Inc., USA). We use OPMs instead of conventional SQUID-MEG system due to the two factors. First of all, with OPMs we are able to register both normal and tangential components of magnetic induction vectors which provides a natural room for comparison of the two technologies capable of measuring the same signals. Secondly, the sensor-scalp distances in the OPM case are comparable to those in the YIGM case. The experimental setup and the scheme of positioning the OPMs on the scalp are depicted in Fig. 2 a) and b) correspondingly We registered alpha-wave activity in the supine position, see Fig.2, a). The recording lasted for 30 seconds in the eyes-open condition and for the next 30 seconds in the eyes-closed condition. This experiment was repeated twice with OPMs calibrated to register normal and tangential components of the magnetic field. As a result of this experiment we obtained individual locations corresponding to the maximal magnitude of the occipital alpha-rhythm in the eyes-closed condition. To do so we used the power in the 8-13 Hz band of the spectrum the OPM registered signals. Since we currently have only a single YIGM sensor we used this individually determined location with top alpha band power to perform our YIGM based measurements described next.

**Figure 2.**
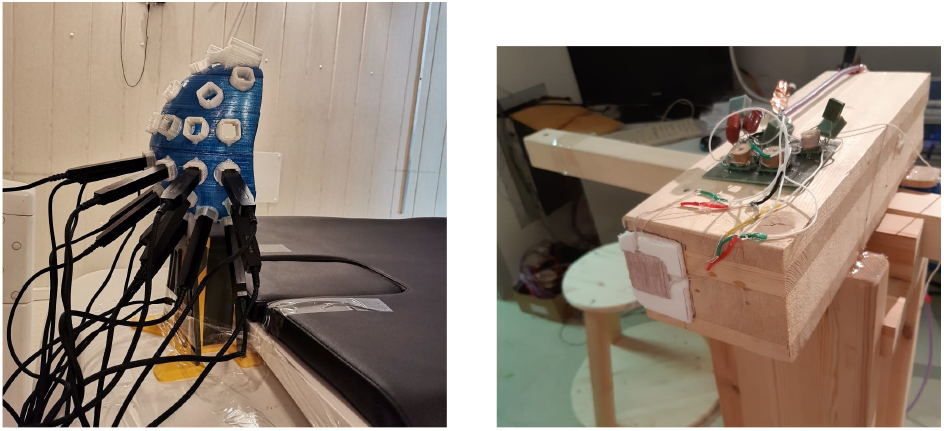
The OP-MEG system used to find locations with high magnitude of alpha waves (a); the YIGM-MEG (1-channel) for alpha-rhythm registration (b).

#### 3.6 The YIG-MEG alpha activity registration experiment

This is the main experiment dedicated to registration of alpha waves with a solid-state YIGM sensor. The experimental setup we used is depicted in Fig. 2, b). A subject is seated on a wooden chair in front of the sensor so that the YIGM sensor is as close as possible to the individually determined location over the occipital lobe corresponding to the maximal alpha power, (see Fig.5). As it was mentioned above (see section 2) YIGM has two sensitive axes (horizontal and vertical) and therefore can detect both tangential and radial field components, depending on mutual position of the head and the sensor (see Fig.3).

For the purposes of this paper, the vertical channel was not recorded due to the high vertical magnetic noise component in our MSR. In order to detect tangential field, the subjects were asked to sit with their head surface tangential to the plane of the YIGM sensor, see Fig. 3, a). In case of the normal field detection, the subjects were asked to rotate 90 degrees with the chair and move to a spot so that the sensor’s plane is orthogonal to the local head surface (Fig. 3, b). Similarly to the OPM-based study we recorded each subject for 30 seconds with eyes-open and then eyes-closed conditions.

**Figure 3.**
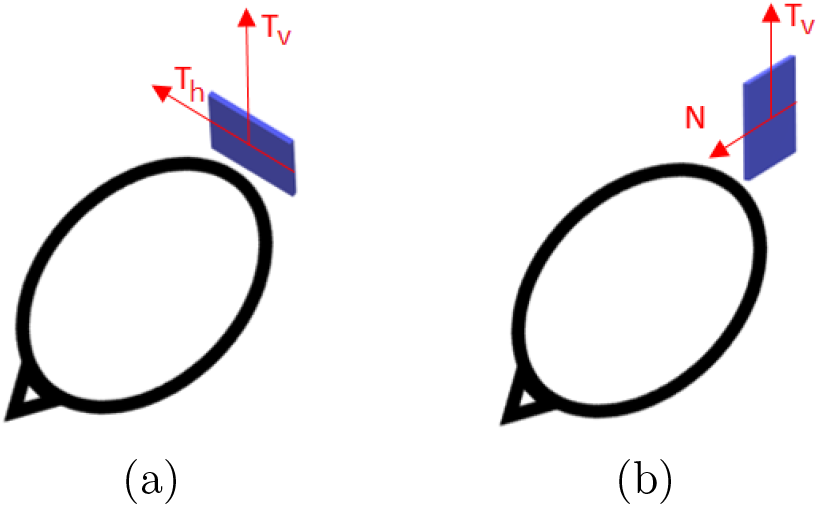
YIGM sensitive axes with respect to mutual head-sensor location: tangential displacement (a); normal displacement (d)

#### 3.7 Analysis of experimental data

Analysis of the obtained data (both OPM and YIGM data) has been performed using Python 3.7 and its modules (SciPy, NumPy, PyPlot, etc.). We used Welch’s method to compute the amplitude spectral density (ASD) of our signals [26]. We employed a 1.5 second long Hamming window and 0.5 seconds overlap to achieve a tradeoff between frequency resolution and variance in our ASD estimates. For filtering our data we used classical one-pass order 3 IIR Butterworth bandpass filters [27] as implemented in SciPy.signal module with cutoff frequencies 9-12 Hz which was selected based on the visual analysis of the individual ASDs.

## 4 Results

### 4.1 YIGM calibration and empty room study

The results of calibrated signal generated by test coil inside the permalloy screen is shown in Fig. 4, a).

**Figure 4.**
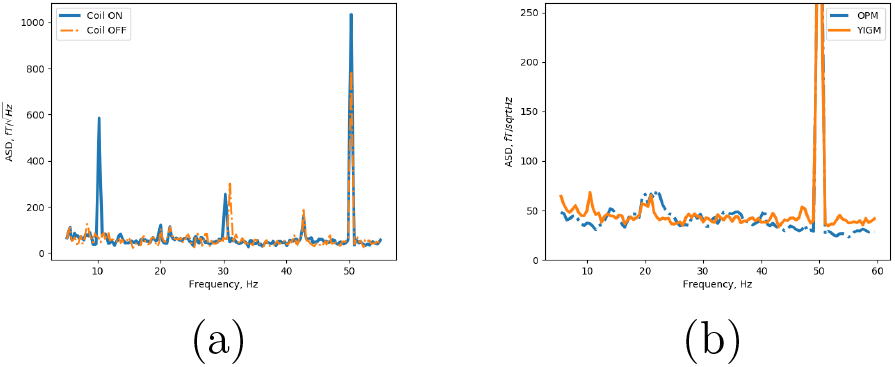
(a) Spectrum of the YIGM-registered signal with the test coil turned on (blue), and turn off (orange) in the empty MSR; (b) Spectrum of the signal registered at the same spatial location in the empty room with YIGM (orange) and with OPM QZFM by QuSpin (blue)

Fig. 4, b) shows the spectrograms of the signal registered in empty MSR by both YIGM and OPM QZFM. Both measurements were conducted at the same location in the MSR. One can see the common peak of the industrial frequency at 50 Hz, and temperate noise at the neighborhood of 20 Hz. Since we see this noise with different-kind of sensors, we assume it to be an outer magnetic contamination presented in MSR, and not a sensor noise. Moreover, this frequency is out of scope of the current article, and thus we do not consider it in details.

Analysis of figure 4 b allows us to conclude that the maximum noise in the neighbourhood of 10 Hz does not exceed 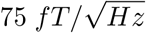. Taking into account that at a 1 cm distance from the skull the expected value of the signal produced by the alpha rhythm is on the order of hundred fT, we can conclude the noise level is acceptable for main our eperiment. In the subsequent presentation for clarity we will focus on the spectra within (5, 18) Hz range.

### 4.2 The OP-MEG alpha activity registration experiment

The purpose of the current experiment was to register alpha waves in all three subjects, and localize the areas on the scalp with highest magnitude of magnetic induction vector field. Such locations for each subject are shown on Fig. 5 in the top panel. The sensors detecting the highest amplitude of signal are highlighted with green color. Corresponding spectrograms are presented in the right panel of the Fig. 5. In the middle panel we show the power spectral density (ASD) of signal obtained using the calibration of OPMs to register normal component of the magnetic field (further - normal calibration). Bottom panel of the figure shows the ASD of the tangential component (tangential calibration).

**Figure 5.**
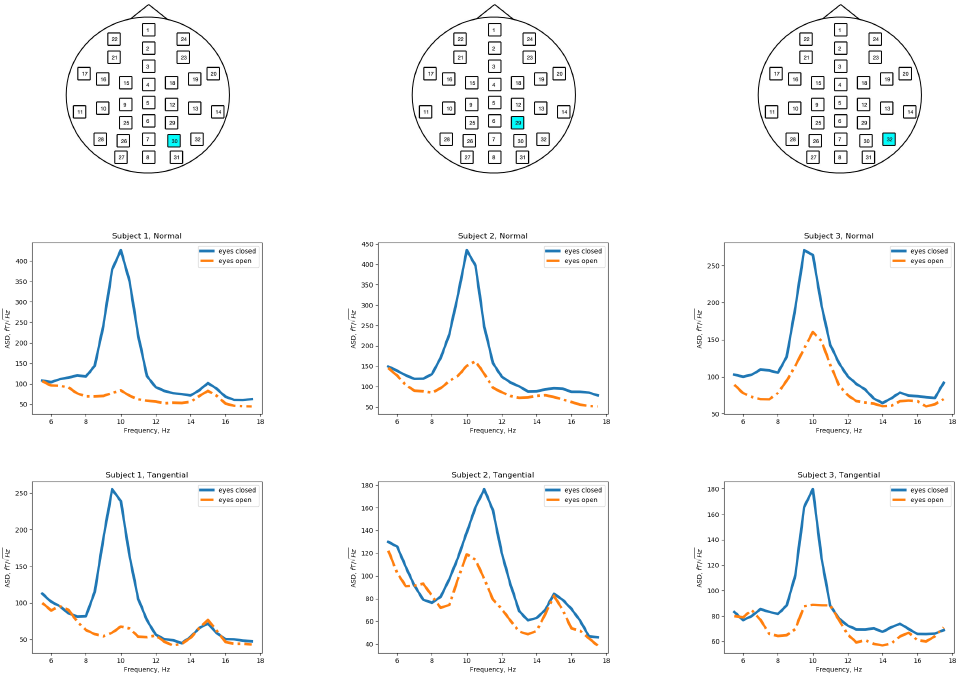
Locations of OPMs for OP-MEG alpha waves registration (top panel); middle and bottom panels - spectra of the signal measured in normal and tangential way, respectively. The spectra correspond to the marked OPMs - OPMs that registered the highest magnitudes of alpha wave.

### 4.3 The YIG-MEG alpha activity registration experiment

This experiment represents the aim of the current study: registration of alpha waves with YIGM.

For this purpose, we place the YIGM into locations found during OP-MEG experiment and depicted on Fig. 5. Changing the position of each subject, we registered normal and tangential components of the magnetic induction vector expecting to obtain alpha rhythm amplitudes comparable with those registered with OPMs.

Fig.6 depicts spectra of the obtained signal for each subject of the research. Spectra of the normal component are presented with the top panel of the Fig. 6, while the bottom panel presents the tangential components.

**Figure 6.**
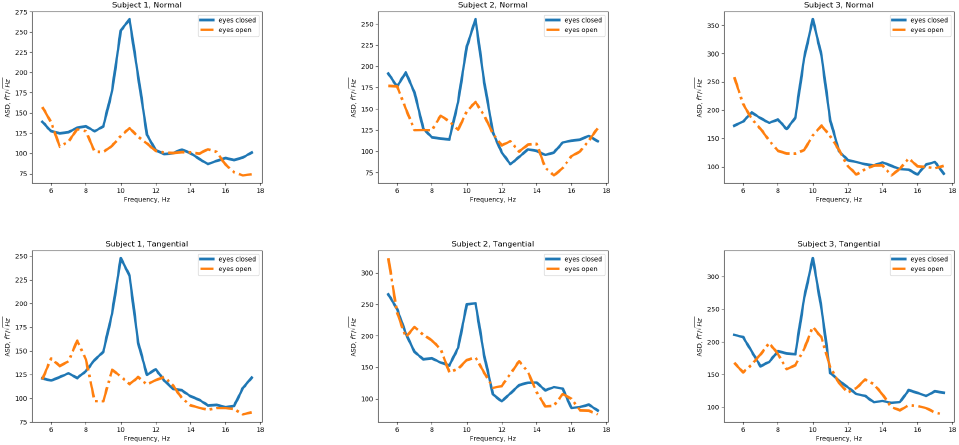
Spectra of YIGM signal measured in all subjects: top panel - normal component of the field registered along *N* sensitive axis of YIGM; bottom panel - tangential component of the field registered along *T*_*h*_ sensitive axis.

## 5 Discussion

As it can be seen from the spectra above (Fig. 5 and Fig. 6), frequency of alpha waves registered in all subjects lies between 9 and 11 Hz, which conforms well with the expected range for subjects of 25-35 y.o. tested in our experiments. In order to isolate the alpha-band oscillation in our data, we filtered the obtained YIGM signals in the (9, 11) Hz band, and superimposed the segments of this data registered during the eyes-open and eyes-closed conditions on a single graph as shown in Fig. 7, a), b), and c) for the three subjects. The blue line represents the closed-eyes signal, while the orange one related to the open-eyes state. We have also visualized our observations in a way similar to that used by D.Cohen in his 1972 paper heralding the use of SQUIDS for human alpha registration. In Fig. 7, d) and e) we show the normal component of the YIGM measured activity around the the moment of closing eyes. One can see the whole experiment duration is close to 1 minute, and subject was asked to close their eyes after 30 seconds from the moment the experiment started. The orange line on that graphs represents the envelope obtained as the absolute value of the band-filtered signal Hilbert transform.

**Figure 7.**
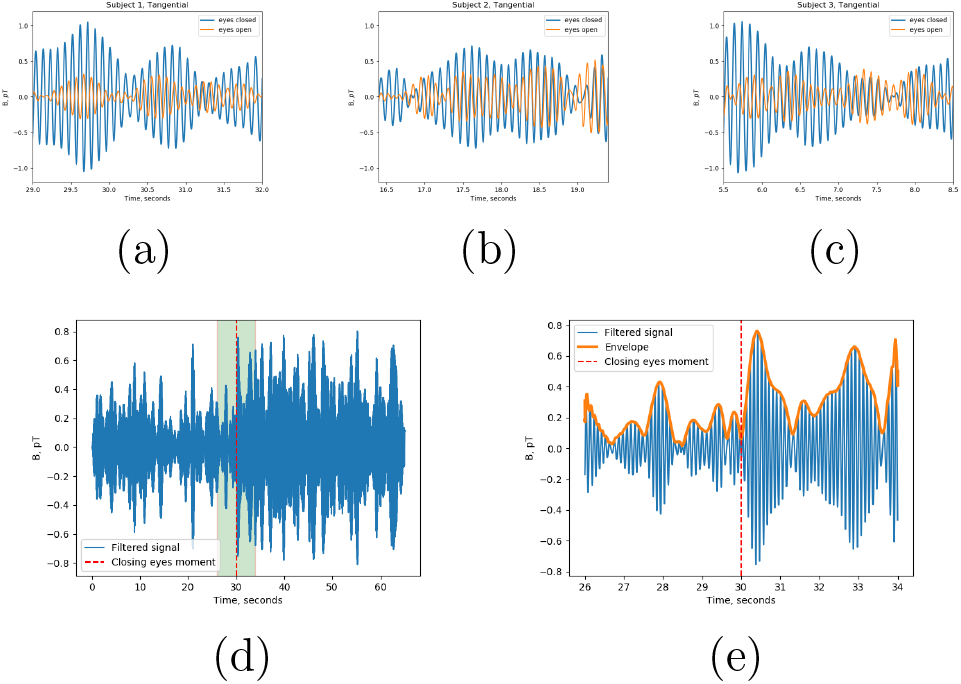
Signal of YIGM obtained during the experiment of registration the tangential component *T*_*h*_ filtered in band (9, 11) Hz. (a), (b), (c) - comparison of signal amplitude in band while eyes are open/closed for each subject (blue line represents closed eyes). (d) and (e) the whole signal filtered in band (9, 11) Hz and its part bounded with green box around the moment of closing the eyes for subject 1, respectively.

Taking a closer look at the ASDs shown in Fig. 5 and Fig. 6, we can compare qualitatively the alpha rhythm registrations obtained with OPM and YIGM. The observed ASD magnitudes appear to be comparable for both sensor types. Interestingly, the second and the third subjects demonstrate more pronounced tangential component of the alpha-rhythm when measured with YIGM as compared to the OPM measurements, showing comparatively low tangential magnitude of alpha wave (see Figures 5 and 6). This observation can be explained by the assessment of distances between the sensors and the hypothesized cortical alpha rhythm generator. The OPM measurements of the normal and tangential components are characterized by one and the same distance to the vapor cell regardless of the sensitive axis used. On the other hand, the distance between scalp and the sensitive element in the YIGM case depends on the mutual arrangement of the head and the sensor, see Figure 3 c and d. During the registration of the normal component, the YIGM is oriented so that its edge points towards the subject’s head as shown in 3.d which increases the mean distance from the brain source to the sensitive element of YIGM as compared to the sensor location when the tangential component is registered, see 3.c. This leads to the observed increase in the magnitude of the tangential components as measured by YIGM sensors. Using an improved setup in our follow-up studies we are planning to more closely investigate this phenomenon which may have serious implications for the future whole head MEG system design.

The design of a multi-channel YIG-MEG system is another major undertaking that we plan for the nearest future. This ambitious work will require producing a number of YIGMs with similar properties, development of the sensor fixation devices and a head-sensor co-registration methodology as well as a detailed exploration of the extent to which the closely located sensors will influence each other’s measurements.

Reassuringly, based on our previous studies, we can expect low level of cross-talks between the sensors. The YIGM considered in this publication has previously been used within the framework of bio-magnetic measurements: detailed studies of the vector distribution of the magnetic field created by the human heart [28, 29], as well as small animals [30] were carried out. Additionally, as a part of the magnetocardiography device development, a three-channel system with three simultaneously operating sensors has been built.

Generally, the interaction of YIGM occurs in two scenarios: 1) interaction of the pump field 2) interaction of the response fields caused by the measured field (second harmonic). The interaction of the pumping fields of the sensors can be minimized by using a coherent (or one) sinusoidal pumping signal source. Also, it is desirable that the rotation of the vectors of neighboring sensors coincide. Small additive changes in the pump fields caused by neighboring sensors can be compensated by changing the pump field in each sensor separately. In fact, these changes are insignificant and their correction was not required when using three sensors simultaneously [29]. The interaction of the response fields is leveled by the feedback system. Since the field is measured by the zero method, i.e. the feedback signal reduces the amplitude of the measured effect to the noise level, so such interaction is not expected and was not detected in the experiment.

We can consider various schemes of positioning YIGM sensors around the human head. One possible way is to simply cover the head surface with tangentially oriented sensor planes resulting into 40-50 vector magnetometers registering tangential field components. To register the radial component the magnetometers can be oriented perpendicularly to the head surface. Sensors located both tangentially and radially can also be combined. It is also possible to construct planar gradiometers based on the measurements of the pairs of nearby magnetometers. Since it is possible to further miniaturize the YIGM sensors [17] the number of sensors in the final system may be increased. The choice of the exact configuration needs to be based on optimizing specific formal criterion such as for example information transfer coefficient which is going to be addressed in our upcoming reports.

## 6 Conclusion

In this work we demonstrated the possibility of the application of a solid state yittrium-iron garnet magnetometer to register brain activity according to the magnetoencephalograhic principle. To show this we designed an experiment for alpha rhythm registration and validated our YIGM measurements with those obtained with commercially available optically pumped magetometers (OPMs).

It is important to emphasize that the YIGM sensor used in this experiment is the first solidstate magnetic field sensor with such a low noise level. The solid-state sensors are in principle, easier to integrate and maintain. The device operates at room temperature, which, first of all, makes it cheaper to maintain than SQUID, and, secondly, allows for further reduction of the distance between the sensor and the brain source. The latter obviously depends on the shape of the sensor: flat shape allows the magnetometer placed right onto the head surface. The sensor implements vector measurement mode by registering magnetic field vector projections onto its surface, which results in measuring two tangential components of the magnetic field simultaneously.

Finally, theoretically it was predicted that this technology has a potential to increase its sensitivity to 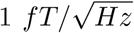 which together with the observations reported above support our idea of developing a multi-channel MEG system based on the fully solid-state magnetic field sensors. Development of such a device will be a seminal step in the MEG hardware and the entire field of fucntional neuroimaging.

## Footnotes

1 The data that support the findings of this study are available from the corresponding author upon reasonable request.

2 The experiment was carried out in accordance with the recommendations of the Declaration of Helsinki and its amendments, and the protocol was approved by the ethics committee of the National Research University Higher School of Economics. All subjects were healthy humans who gave written informed consent in accordance with the Declaration of Helsinki.

## References

[1] D. Cohen, Magnetoencephalography: detection of the brain’s electrical activity with a superconducting magnetometer, Science 175 (4022) (1972) 664–666.

[2] A. Koptelova, R. Bikmullina, M. Medvedovsky, S. Novikova, A. Golovteev, O. Grinenko, M. Korsakova, A. Kozlova, N. Arkhipova, A. Vorobyev, et al., Ictal and interictal meg in pediatric patients with tuberous sclerosis and drug resistant epilepsy, Epilepsy research 140 (2018) 162–165.

[3] P. K. Mandal, A. Banerjee, M. Tripathi, A. Sharma, A comprehensive review of magnetoencephalography (meg) studies for brain functionality in healthy aging and alzheimer’s disease (ad), Frontiers in Computational Neuroscience 12 (2018) 60.

[4] L. I. Boon, V. J. Geraedts, A. Hillebrand, M. R. Tannemaat, M. F. Contarino, C. J. Stam, H. W. Berendse, A systematic review of meg-based studies in parkinson’s disease: The motor system and beyond, Human brain mapping 40 (9) (2019) 2827–2848.

[5] R. Hari, R. Salmelin, Magnetoencephalography: from squids to neuroscience: Neuroimge 20th anniversary special edition, Neuroimage 61 (2) (2012) 386–396.

[6] F. L. da Silva, Eeg and meg: relevance to neuroscience, Neuron 80 (5) (2013) 1112–1128.

[7] S. Baillet, Magnetoencephalography for brain electrophysiology and imaging, Nature neuroscience 20 (3) (2017) 327–339.

[8] F. Pulvermüller, Y. Shtyrov, R. Ilmoniemi, Spatiotemporal dynamics of neural language processing: an meg study using minimum-norm current estimates, Neuroimage 20 (2) (2003) 1020–1025.

[9] E. Boto, N. Holmes, J. Leggett, G. Roberts, V. Shah, S. S. Meyer, L. D. Muñoz, K. J. Mullinger, T. M. Tierney, S. Bestmann, et al., Moving magnetoencephalography towards real-world applications with a wearable system, Nature 555 (7698) (2018) 657–661.

[10] A. Borna, T. R. Carter, J. D. Goldberg, A. P. Colombo, Y.-Y. Jau, C. Berry, J. McKay, J. Stephen, M. Weisend, P. D. Schwindt, A 20-channel magnetoencephalography system based on optically pumped magnetometers, Physics in Medicine & Biology 62 (23) (2017) 8909.

[11] T. M. Tierney, N. Holmes, S. S. Meyer, E. Boto, G. Roberts, J. Leggett, S. Buck, L. Duque-Muñoz, V. Litvak, S. Bestmann, et al., Cognitive neuroscience using wearable magnetometer arrays: Non-invasive assessment of language function, NeuroImage 181 (2018) 513–520.

[12] C.-H. Lin, T. M. Tierney, N. Holmes, E. Boto, J. Leggett, S. Bestmann, R. Bowtell, M. J. Brookes, G. R. Barnes, R. C. Miall, Using optically pumped magnetometers to measure magnetoencephalographic signals in the human cerebellum, The Journal of physiology 597 (16) (2019) 4309–4324.

[13] E. Boto, S. S. Meyer, V. Shah, O. Alem, S. Knappe, P. Kruger, T. M. Fromhold, M. Lim, P. M. Glover, P. G. Morris, et al., A new generation of magnetoencephalography: Room temperature measurements using optically-pumped magnetometers, NeuroImage 149 (2017) 404–414.

[14] A. Borna, T. R. Carter, A. P. Colombo, Y.-Y. Jau, J. McKay, M. Weisend, S. Taulu, J. M. Stephen, P. D. Schwindt, Non-invasive functional-brain-imaging with an opm-based magnetoencephalography system, Plos one 15 (1) (2020) e0227684.

[15] R. M. Hill, E. Boto, M. Rea, N. Holmes, J. Leggett, L. A. Coles, M. Papastavrou, S. K. Everton, B. A. Hunt, D. Sims, J. Osborne, V. Shah, R. Bowtell, M. J. Brookes, Multi-channel whole-head opm-meg: Helmet design and a comparison with a conventional system, NeuroImage 219 (2020) 116995. doi:https://doi.org/10.1016/j.neuroimage.2020.116995.

[16] P. M. Vetoshko, M. V. Valeiko, P. I. Nikitin, Epitaxial yttrium iron garnet film as an active medium of an even-harmonic magnetic field transducer, Sensors and Actuators A: Physical 106 (1-3) (2003) 270–273.

[17] P. Vetoshko, Remagnetization of iron-garnet films by coherent rotation for sensitive elements of magnetic sensors (in russian), Ph.D. thesis, M.N. Mikheev Institute of Metal Physics of the Ural Branch of the Russian Academy of Sciences (IMP UB RAS) (2017).

[18] J. Osborne, J. Orton, O. Alem, V. Shah, Fully integrated standalone zero field optically pumped magnetometer for biomagnetism, in: Steep Dispersion Engineering and Opto-Atomic Precision Metrology XI, Vol. 10548, International Society for Optics and Photonics, 2018, p. 105481G.

[19] T. Starner, Human-powered wearable computing, IBM systems Journal 35 (3.4) (1996) 618–629.

[20] M. Steriade, Corticothalamic resonance, states of vigilance and mentation, Neuroscience 101 (2) (2000) 243–276.

[21] M. Steriade, Impact of network activities on neuronal properties in corticothalamic systems, Journal of neurophysiology 86 (1) (2001) 1–39.

[22] P. Roland, Cortical organization of voluntary behavior in man., Human neurobiology 4 (3) (1985) 155–167.

[23] G. Buzsaki, Rhythms of the Brain, Oxford University Press, 2006.

[24] T. Ergenoglu, T. Demiralp, Z. Bayraktaroglu, M. Ergen, H. Beydagi, Y. Uresin, Alpha rhythm of the eeg modulates visual detection performance in humans, Cognitive Brain Research 20 (3) (2004) 376–383.

[25] F. H. D. Silva, Encyclopedia of the Human Brain, Elsevier, 2002, Ch. Electrical potentials, pp. 147–167.

[26] P. Welch, The use of fast fourier transform for the estimation of power spectra: A method based on time averaging over short, modified periodograms, IEEE Transactions on Audio and Electroacoustics 15 (2) (1967) 70–73. doi:10.1109/TAU.1967.1161901.

[27] G. Bianchi, R. Sorrentino, Electronic filter simulation & design, McGraw Hill Professional, 2007.

[28] N. Gusev, P. Vetoshko, A. Kuzmichev, D. Chepurnova, E. Samoilova, A. Zvezdin, A. Korotaeva, V. Belotelov, Ultra-sensitive vector magnetometer for magnetocardiographic mapping., Biomedical Engineering 51 (3) (2017).

[29] P. Vetoshko, N. Gusev, D. Chepurnova, E. Samoilova, I. Syvorotka, I. Syvorotka, A. Zvezdin, A. Korotaeva, V. Belotelov, Fluxgate magnetic field sensor based on yttrium iron garnet films for magnetocardiography investigations, Technical Physics Letters 42 (8) (2016) 860–864.

[30] P. Vetoshko, N. Gusev, D. Chepurnova, E. Samoilova, A. Zvezdin, A. Korotaeva, V. Belotelov, Rat magnetocardiography using a flux-gate sensor based on iron garnet films., Biomedical Engineering 50 (4) (2016).

